# LEDGF and HDGF2 relieve the nucleosome-induced barrier to transcription

**DOI:** 10.1101/500926

**Authors:** Gary LeRoy, Ozgur Oksuz, Nicolas Descostes, Yuki Aoi, Rais A. Ganai, Havva Ortabozkoyun Kara, Jia-Ray Yu, Chul-Hwan Lee, James Stafford, Ali Shilatifard, Danny Reinberg

**Author notes:** These authors contributed equally to this work (Gary LeRoy, Ozgur Oksuz, Nicolas Descostes).

## Abstract

FACT (<u>fa</u>cilitates <u>c</u>hromatin <u>t</u>ranscription) is a protein complex that allows RNAPII to overcome the nucleosome-induced barrier to transcription. While abundant in undifferentiated cells and many cancers, FACT is not abundant or is absent in most tissues. Therefore, we screened for additional proteins that might replace FACT upon differentiation. Here we report the identification of two such proteins, LEDGF and HDGF2, each containing two HMGA-like AT-hooks and a PWWP methyl-lysine reading domain known to bind to H3K36me2 and H3K36me3. LEDGF and HDGF2 localize with H3K36me2/3 at genomic regions containing active genes, usually adjacent to H3K27me3 domains. In myoblasts, where FACT expression is low, LEDGF and HDGF2 are enriched on most active genes. Upon differentiation to myotubes, LEDGF levels decrease across the genome while HDGF2 levels are maintained. Moreover, HDGF2 is recruited to the majority of myotube up-regulated genes and is required for their proper expression. HDGF2 knockout myoblasts exhibit an accumulation of paused RNAPII proximal to the first nucleosome within the transcribed region of many HDGF2 target genes, indicating a defect in early elongation. We propose that LEDGF and HDGF2 substitute for FACT in differentiated tissues and that their distribution on the genome helps maintain transcriptional programs unique to particular cell types.

**One Sentence Summary:** Chromatin bound LEDGF and HDGF2 proteins allow RNAPII to overcome the nucleosome-induced block to transcription.

## Main Text

RNA polymerase II (RNAPII) transcription is regulated at the level of initiation, promoter escape, pause release and elongation (*1, 2*). After pause release, RNAPII must overcome a nucleosome-induced barrier to transcription (*3, 4*). Two decades ago, we identified FACT (<u>fa</u>cilitates <u>c</u>hromatin <u>t</u>ranscription) as a protein complex composed of SPT16 and SSRP1 that alleviates this nucleosome-induced barrier to transcription (*5, 6*). More recently, we mapped the genomic binding of FACT in stem cells using ChIP-seq with an SPT16 antibody and found that FACT occupancy varies, being associated at high levels with only a subset of active genes (**Fig. 1A**). In addition, we and others have found that FACT is only expressed at high levels in some progenitor and transformed cells (**Fig. 1B**) (*7, 8*). These observations imply the existence of alternative FACT-like chaperones in differentiated cells. We first hypothesized that the BET family proteins (Brd2, Brd3 and Brd4), which also possess FACT-like activity *in vitro*, might replace FACT at these genes (*9, 10*). However, BET proteins predominately localize with acetylated nucleosomes proximal to the transcription start site (TSS) of genes, while a novel chaperone capable of fulfilling the function of FACT should localize in gene bodies (*10, 11*).

**Figure 1.**
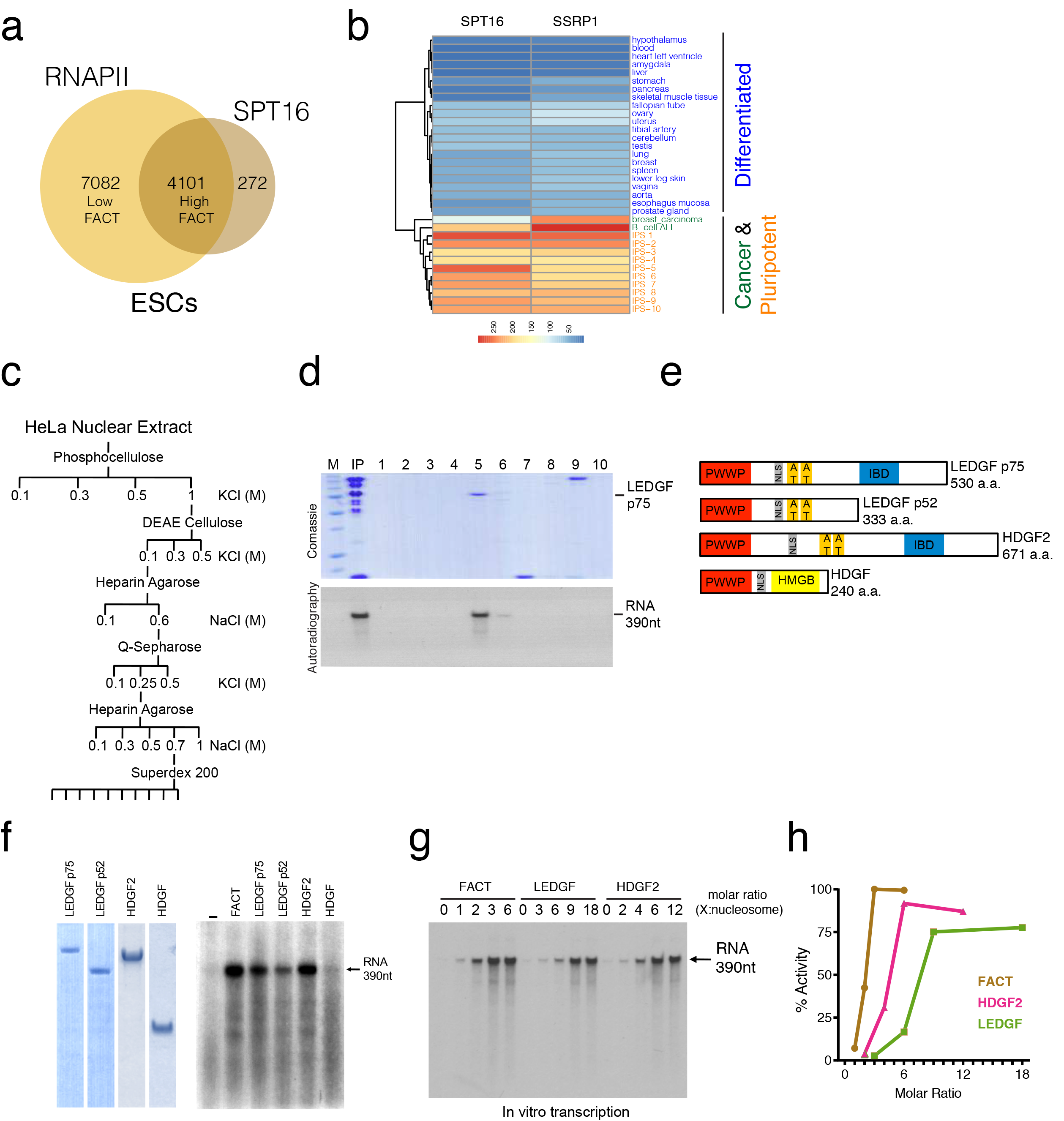
Biochemical screen identifies LEDGF and HDGF2 as FACT-like factors. **A.** Venn diagram showing occupancy of RNAPII and SPT16 (FACT) on active genes (at least 5 transcripts) in mouse stem cells. Hypergeometric test p-value = 0. **B.** Expression of FACT subunits, SPT16 and SSRP1 in different cell types. Heatmaps were created using normalized RNA-seq data and ordered by hierarchical clustering. **C.** Schematic depicting the chromatographic steps of the biochemical screen used to obtain a purified fraction containing a novel FACT-like activity. **D.** Top panel, fractions from the Superdex-200 step were separated by SDS-PAGE, stained with Coomassie Blue, and analyzed by MS for protein identification. Bottom panel, fractions from the Superdex-200 step analyzed using the *in vitro* chromatin transcription assay. **E.** Schematic of the predicted domain structure of the LEDGF/HDGF2 family of proteins. PWWP = methyl-lysine binding domain, NLS = nuclear localization sequence, AT = AT hook domain, HMGB = High mobility group box domain and IBD = Integrase binding domain. **F.** Left panel, the highly purified recombinant versions of the LEDGF/HDGF2 family proteins were separated by SDS-PAGE and stained with Coomassie Blue. Right panel, highly purified recombinant versions of the LEDGF/HDGF2 family proteins analyzed using the *in vitro* chromatin transcription assay. **G.** Titration of FACT, LEDGF and HDGF2 in defined RNAPII transcription assays with nucleosomal templates. Molar ratio of protein (X):nucleosome in assays is indicated on top. **H.** Graph of transcription quantified from (H). Y-Axis is the relative activities quantified with Image J software. X-Axis is the molar ratio of protein (X):nucleosome in assays.\

Exploiting a similar biochemical strategy that first led us to identify FACT, we fractionated HeLa cell nuclear extract and identified a fraction that was depleted of FACT, BET proteins and Nucleolin (RNA Polymerase I-specific FACT-like chaperone), yet able to support transcription through nucleosomes (**fig. S1A & S1B**) (*5, 9, 10, 12*). Following further chromatographic fractionation and mass spectrometry we identified LEDGF long isoform (p75) (also known as PSIP1) (**Figs. 1C-D**) (*13, 14*). Semi-quantitative proteomics and RNA-seq data suggests that this protein as well as its family member, HDGF2, are relatively abundant and expressed in most tissues (not shown), unlike the restricted expression of FACT. LEDGF and HDGF2 each contain a methyl-lysine reading PWWP domain that has been shown to recognize H3K36me2/3, two HMGA-like AT hooks (similar to Nucleolin) and an Integrase Binding Domain (IBD) (**Fig. 1E**) (*15–19*). The short isoform of LEDGF (p52) and another PWWP containing protein, HDGF, lack the IBD and the latter contains an HMGB domain instead of HMGA-like AT hooks (*20*). LEDGF and HDGF2 have generated considerable interest given their requirement for lentiviral integration, which favors integration into the body of transcribed genes by concurrently binding directly to lentiviral integrase and H3K36me2/3 in the host cell chromatin (*16, 21*). Interestingly, retroviruses which favor integration near the TSS of transcribed genes utilize BET proteins in an analogous manner (*22, 23*). LEDGF and HDGF2 share another similarity with BET proteins in that they remain bound to mitotic chromosomes suggesting they might contribute to transcriptional memory (*24–26*).

To validate the FACT-like activity of LEDGF and its related proteins, we generated and tested purified recombinant versions of these proteins in an *in vitro* reconstituted chromatin transcription assay (**Fig. 1F**). As depicted, both isoforms of LEDGF (p75 and p52) as well as HDGF2 allow RNAPII to transcribe through nucleosomes in a manner similar to FACT. HDGF which lacks the HMGA-like AT hooks did not substitute for FACT, suggesting that the FACT-like activity of these proteins probably lies within the region that contains the AT hooks (**Fig. 1E**) (*20*). Increasing amounts of FACT, LEDGF or HDGF2 led to a linear increase in transcription activity such that at least two or more molecules of LEDGF and HDGF2 per nucleosome appeared to be required for efficient transcription in the assay (**Figs. 1G-I**).

To investigate the functions of LEDGF and HDGF2 in RNAPII transcription *in vivo*, we first asked whether their binding correlates with regions having RNAPII occupancy and variable levels of FACT initially in 293T cells (**Fig. 2B**). Of note, all three factors; FACT, LEDGF and HDGF2 are highly expressed in 293T cells unlike more differentiated cells (**Fig. 1B & 2A**). Thus, we performed native ChIP-seq using FLAG antibody in 293T cells stably expressing FLAG-LEDGF and FLAG-HDGF2, and compared their occupancy with SPT16, H3K36me2, H3K36me3 and H3K27me3 on RNAPII bound loci (**Figs 2C & S2A**). We also performed ChIP-seq with LEDGF antibody in 293T cells and observed a similar binding pattern to that of FLAG-LEDGF, validating the antibody to be used in later experiments (not shown). After selecting RNAPII bound loci, K-mean clustering segregated the loci into six clusters based on high, medium and low levels of either FACT or LEDGF/HDGF2 respectively (fig. S2A). SPT16 most closely correlated with high levels of RNAPII binding in accordance with its high levels in 293T cells (**Fig. 2A**). Although some of the genes having high levels of RNAPII were enriched with all three factors (SPT16, LEDGF and HDGF2), many genes have low levels SPT16 (low FACT) and instead were enriched with LEDGF and HDGF2 (**Fig. 2C and S2A**). These findings suggest that LEDGF and HDGF2 may facilitate transcription of a subset of genes that have low levels of FACT.

**Figure 2.**
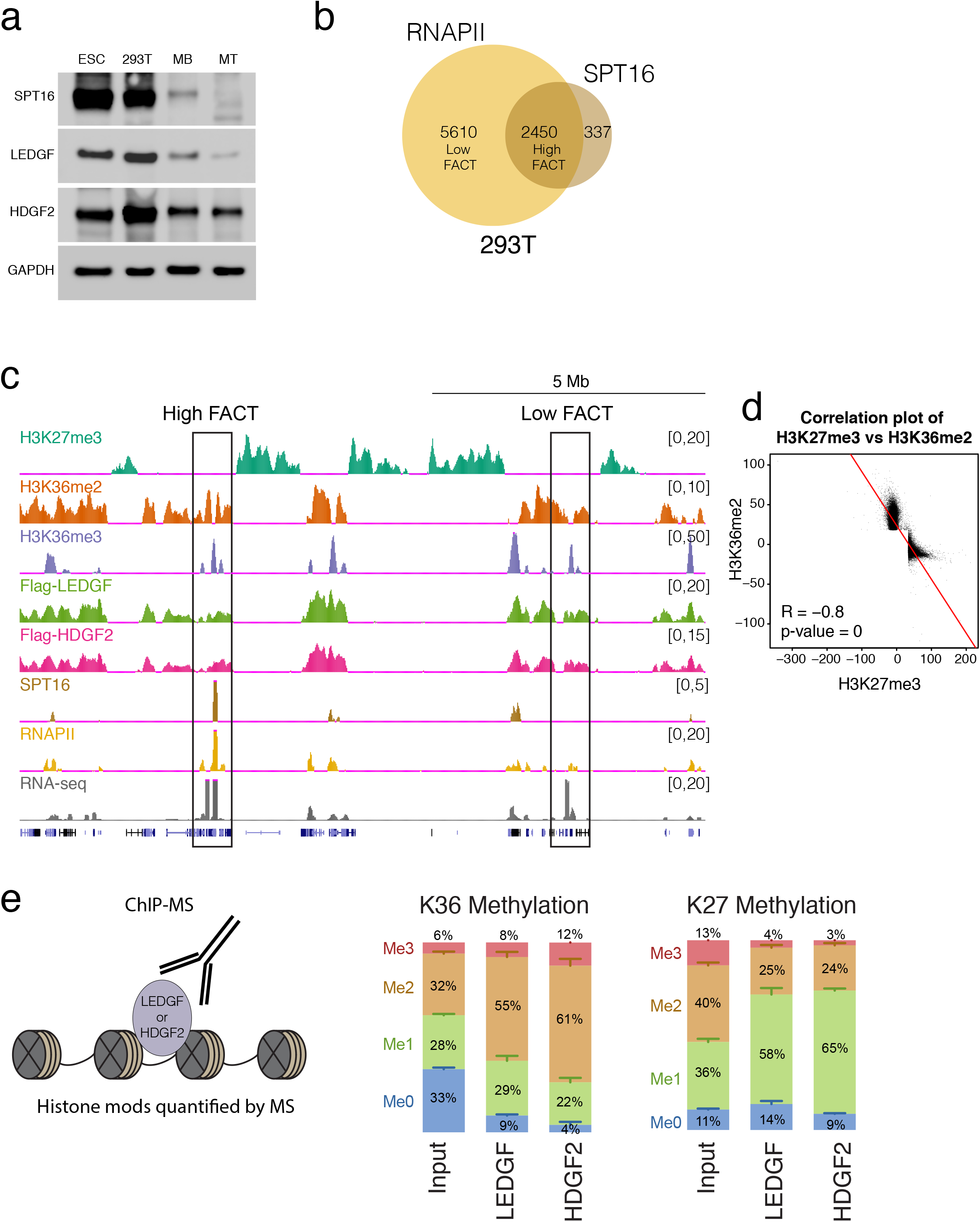
LEDGF and HDGF2 bind to clusters of active chromatin. **A.** Western blots performed with whole cell protein extracts from mESCs, 293T, myoblasts (MB) and myotubes (MT) using the antibodies indicated. **B.** Venn diagram showing occupancy of RNAPII and SPT16 (FACT) on active genes (at least 5 transcripts) in human 293Tcells. Hypergeometric test p-value = 0. **C.** ChIP-seq genomic tracks of the indicated factors at a (~14 MB) region of chromosome 6 with corresponding RNA-seq. **D.** Spearman correlation plot showing the opposition of H3K27me3 and H3K36me2. Union of the Top 20% of peaks were selected. **E.** Left panel, cartoon depicting LEDGF or HDGF2 purified chromatin used to quantify histone modifications by Mass Spectrometry (MS). Right panel, histone H3K27 and H3K36 methylations quantified by Mass Spectrometry from the input (293T whole genome chromatin), FLAG-LEDGF and FLAG-HDGF2 ChIPs.

Our detailed bioinformatics analysis revealed that SPT16 binding closely correlated with highly expressed housekeeping genes, whereas LEDGF and HDGF2 were more enriched on genes greater in length that exhibit a higher frequency of RNAPII pausing (figs. S2B-D). In contrast to SPT16, we observed a striking correlation between the localization of LEDGF and HDGF2 with H3K36me2 (**Figs. 2C & S2A**), consistent with their ability to bind both H3K36me2 and H3K36me3 through their PWWP domains (**15, 18**). For example, similar to that of H3K36me2 domains, ChIP-seq profiles for LEDGF and HDGF2 on an approximately 12 Mb track of chromosome 6 showed that these factors decorate broad domains which include clusters of active genes that are opposed by domains demarked by H3K27me3 (**Fig 2C**). At this megabase level of visualization it is apparent that SPT16 binding is variable, not equally enriched on all clusters of actively transcribed genes and when present, is more restricted to only transcribed genes similar to H3K36me3 and RNAPII. A more detailed bioinformatics analysis of these data utilizing Markov modeling reinforces this partitioning at the genomic level (**fig. S3**). We also found an unparalleled opposition between H3K27me3 and H3K36me2 across the genome (**Fig. 2D**). These modifications and their opposition probably co-evolved together as lower organisms such as *S. Cerevisiae* have neither <u>P</u>olycomb <u>R</u>epressive <u>C</u>omplex <u>2</u> (PRC2), the sole enzyme responsible for H3K27 methylation, nor dedicated H3K36 di-methylase enzymes (*27*). Interestingly, *S. Cerevisiea* does not contain a homolog of LEDGF/HDGF2, though *Drosophila* does, an organism that also has PRC2 and dedicated H3K36 di-methylase enzymes (*28, 29*).

We next characterized the histone modifications associated with the FLAG-LEDGF and FLAG-HDGF2 by quantitative ChIP-MS (*11*). As shown in Figure **2E**, the nucleosomes associated with both LEDGF and HDGF2 were highly enriched in H3K36me2 as compared with whole genome histones (Input). Interestingly, HDGF2 also enriched H3K36me3 suggesting that these proteins have a somewhat different binding preference. The nucleosomes associated with both proteins were depleted of the repressive chromatin associated modifications, H3K27me2 and H3K27me3 (**27**). These results are consistent with our ChIP-seq data and provide more direct evidence that these proteins bind H3K36me2 and H3K36me3 *in vivo* (**Fig. 2C**). Though we were not able to reconstitute the dependency on these modifications *in vitro* using the reconstituted transcription assay with synthetic modification-mimic histones (data not shown), the PWWP domains of LEDGF and HDGF2 were both shown to be required for these proteins to bind chromatin *in vivo* (*26, 30*). This discrepancy is likely due to other domains (i.e., AT-hooks) of the LEDGF and HDGF2 proteins which exhibit DNA and nucleosome binding affinities independent of their PWWP domains and are sufficient to bind unmodified nucleosomes in a defined *in vitro* assay (*15*).

To determine if LEDGF and HDGF2 are recruited to genes upon induction, we performed ChIP-seq in mESCs before and after differentiation to embryoid bodies using knockout-validated LEDGF and HDGF2 antibodies (**fig. S4**). We also performed ChIP-seq for SPT16, RNAPII, H3K27me3, H3K36me2 and H3K36me3. However, in order to quantitatively analyze the ChIP-seq data from these two conditions (ESC vs EB), we performed these and all subsequent experiments with spike-in controls (*31*). Interestingly, in mESCs LEDGF is rather dispersed, being found in large domains usually containing some active genes, whereas HDGF2 exhibits a binding pattern similar to SPT16, being more restricted to the bodies of actively transcribed genes (**Figs. 3A-B**). Upon differentiation (ESC to EB), approximately 881 genes showed a 2-fold (or more) increase in RNAPII binding and HDGF2 was recruited to 40% of these genes (**Fig. 3C**). The ChIP-seq binding patterns of SPT16 revealed that it was also recruited to 58% of these genes and had a 23% overlap with HDGF2 (**Fig. 3C**). Moreover, the opposite trend holds true, as genes with reduced RNAPII (ESC to EB) usually exhibited a decrease in HDGF2 and SPT16 binding. Representative examples of specific genes showing these phenomena are presented in **Figs. 3A** and **3B**. These results indicate that both HDGF2 and FACT follow RNAPII deposition at a subset of genes during mESCs differentiation.

**Figure 3.**
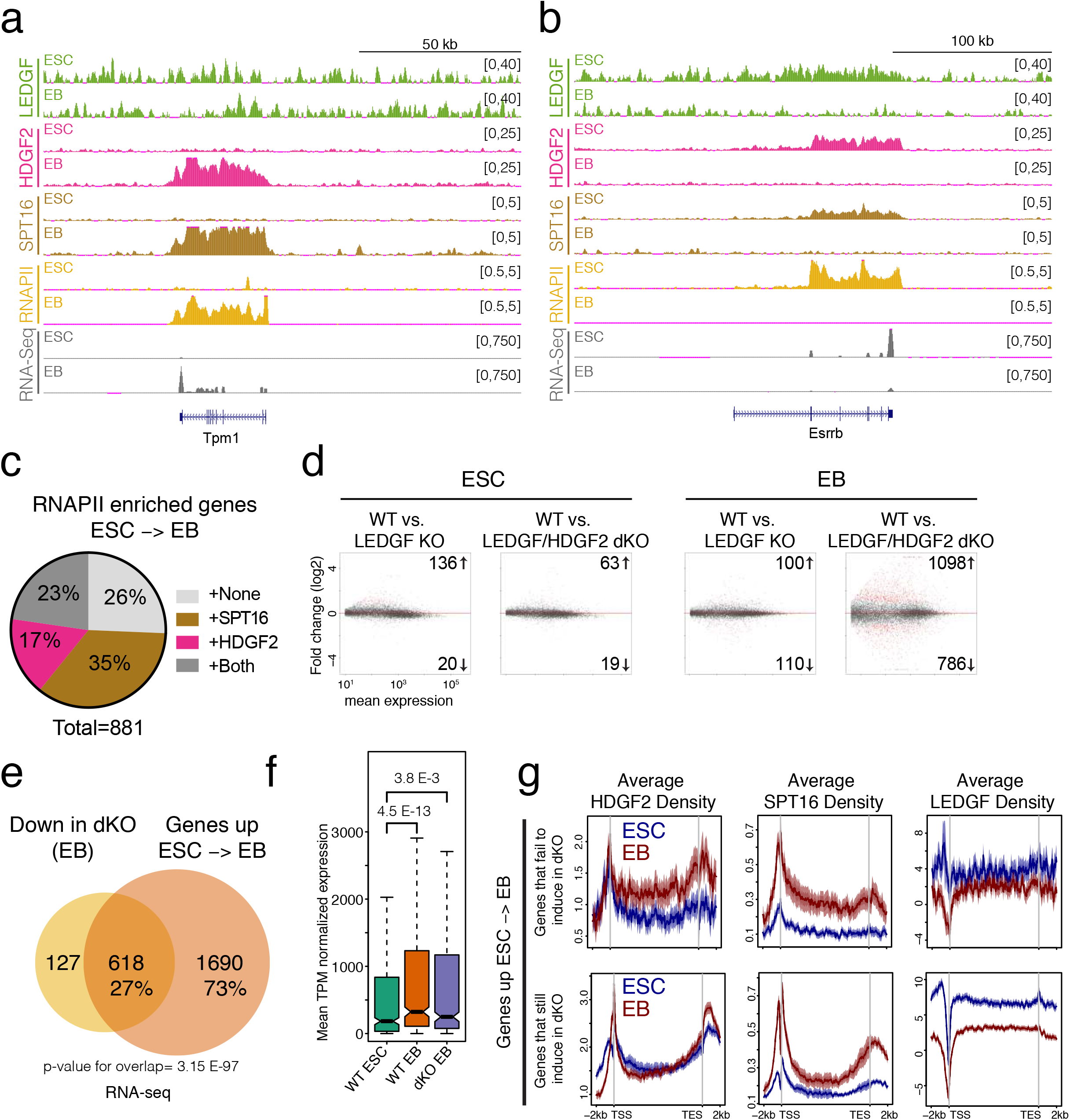
LEDGF and HDGF2 are required for the induction of some genes in stem cells. **A., B.** ChIP-seq tracks for LEDGF, HDGF2, SPT16 and RNAPII with corresponding RNA-seq tracks in ESCs and in EBs at the Tpm1 gene (**A**) and Essrb gene (**B**). **C.** Pie chart showing the percentage of genes exhibiting increased HDGF2 and/or FACT binding on genes with an increase in RNAPII during ESCs differentiation into EBs. **D.** MA plots showing the number of differentially expressed genes (2-fold, BH-corrected p-value < 0.05) in LEDGF KO and LEDGF/HDGF2 dKO in ESCs to EBs differentiation. **E.** Venn diagram depicting the number of up-regulated genes (2-fold) from ESC to EB in WT cells overlaid with the down-regulated genes in the LEDGF/HDGF2 dKO EBs. Of the 2308 genes normally up-regulated in the WT cells (ESC to EB), 618 were found to be down regulated in the dKO EB cells. The overlap with dKO down regulated genes is significant (p-value = 3.15e-97, hypergeometric test). **F.** Boxplots with confidence intervals of expression of the 618 genes selected from **Fig. 3E** at WT ESC, WT EB and dKO EB. Wilcoxon Rank Sum test p-values: WT ESCs vs. WT EBs = 4.5e-13; WT ESCs vs. dKO EBs = 3.8e-13. **G.** Top panels, average density ChIP-seq profiles of HDGF2, SPT16 and LEDGF on the 618 genes that are up-regulated in WT EBs and down regulated in dKO EBs. Bottom panels, average density ChIP-seq profiles of HDGF2, SPT16 and LEDGF on the 4,999 genes identified as not having any change in expression between ESCs and EBs.

In order to validate that LEDGF and HDGF2 participate in transcription we utilized CRISPR/Cas9 technology to knockout (KO) LEDGF alone or together with HDGF2 (LEDGF/HDGF2 dKO) in mESCs. We did not observe a large number of genes whose expression (RNA-seq, 2-fold-change cutoff, BH corrected p-value < 0.05) was affected in either KO cell line compared to WT mESCs (**Fig. 3D**), suggesting that there is a redundancy likely due to high levels of FACT or other unknown factors (**Fig. 2A**). Interestingly, when we differentiate these mESCs into EBs, we observed a significant number of genes (1884 genes) affected in the LEDGF/HDGF2 dKO cells (**Fig. 3D**). However, the changes in the LEDGF KO cells were still minimal, implying a redundancy between LEDGF and HDGF2 at this stage of cellular differentiation.

Given that HDGF2 was recruited to approximately 40% of the genes that increase expression during differentiation of ESCs to EBs (**Fig 3C**), we next analyzed the effect of the LEDGF/HDGF2 dKO on these upregulated genes. Upon differentiation (ESC to EB), 2308 genes exhibited at least a 2-fold increase in expression in WT cells as measured by RNA-seq (**Fig. 3E**). Of these differentially expressed genes, ~27% failed to fully activate in the LEDGF/HDGF2 dKO cells (**Figs. 3E-F**). Average ChIP-seq read density profiles showed a correlation between these genes and an increase in HDGF2 binding (ESC to EB) (**Fig. 3G top left panel**). Conversely, we did not observe this correlation on the genes that were activated to the same extent in dKO cells as in WT cells (**Fig. 3G, bottom left panel**). However, such genes did correlate with an increase in SPT16 binding, possibly indicating redundancy (**Fig 3G, bottom middle panel**). SPT16 binding generally showed an overall increase on most genes that gain expression from ESC to EB, even those genes that failed to properly induce in the dKO cells (**Fig. 3G**). Such redundancy could explain why many of the induction-compromised genes still showed partial activation in the dKO cells (**Fig. 3F**).

To control for the possible redundancy between LEDGF/HDGF2 and FACT, we utilized myoblast (MB) to myotube (MT) cellular differentiation system (*32, 33*) as protein levels of FACT (SPT16 and SSRP1) and LEDGF decrease while HDGF2 remain constant during differentiation into MT (**Fig 4A**). As expected, binding of both LEDGF and HDGF2 on the genome correlated with RNAPII levels as evidenced by average density profiles in MB (**Fig. 4B upper panels & S5A**). However, SPT16 was only detected at low levels on a few highly expressed genes in MB. Upon differentiation into MT, SPT16 was not detected on any genes and LEDGF binding decreased globally while HDGF2 remained enriched on actively transcribed loci and accumulated on genes whose expression was induced (**Figs. 4B lower panels & S6D**).

**Figure 4.**
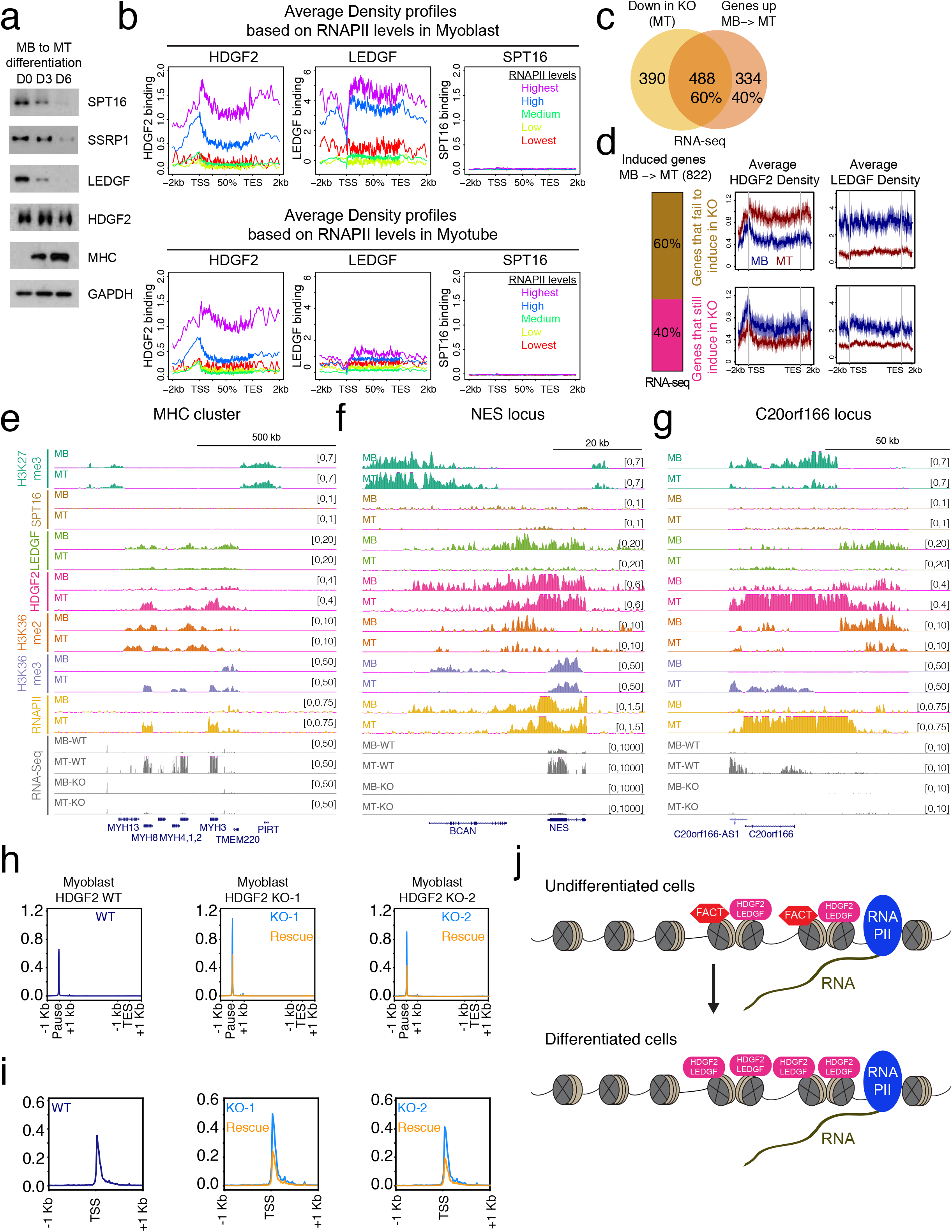
LEDGF and HDGF2 substitute for FACT in differentiated cells. **A.** Western blots of whole cell protein extracts from myoblasts (MB) differentiated to myotubes (MT) obtained from Day 0 (D0), Day 3 (D3) and Day 6 (D6) using the antibodies indicated. **B.** Average density ChIP-seq profiles for HDGF2, LEDGF and SPT16 based on genes exhibiting RNAPII binding in MB (upper panels) and MT (lower panels). Levels based on genes with RNAPII binding, Highest (purple, top 5%), High (blue, top 5-20%), Medium (dark green, 20-80%), Low (bright green, bottom 5-20%) and Lowest (red, bottom 5%). **C.** Venn Diagram depicting the number of genes up-regulated 2-fold in MB to MT cells overlaid with genes down-regulated 2-fold in HDGF2 KO MT cells. **D.** Average density ChIP-seq profiles for HDGF2 and LEDGF in MB and MT cells. Top panels, the 488 genes that are up-regulated in MB to MT and fail to induce in the HDGF2 KO cells. Bottom panels, the 334 genes that still induce in the HDGF2 KO. **E-G.** ChIP-seq tracks for H3K27me3, SPT16 LEDGF, HDGF2, H3K36me3 and RNAPII with the corresponding RNA-seq tracks for MB and MT at the MHC cluster (**E**), the NES locus (**F**) and the C20orf166 locus (**G**). **H-I.** Metagene profiles of PRO-seq RNA data plotted for the top 20% HDGF2 bound genes in WT and HDGF2 KO and HDGF2 KO rescue myoblast cell lines as indicated on the panels. (**H**) Plotted by 1kb-Pause site-1kb-(Scaled gene bodies)-1kb-TES-1kb. (**I**) Plotted with data centered on the TSS (-/+ 1kb). **J.** Schematic depiction showing the replacement of FACT by HDGF2 and/or LEDGF on chromatin during cellular differentiation, when FACT expression is reduced.

As these data indicate that a MB to MT differentiation system is ideal for studying the function of HDGF2, we generated a HDGF2 KO in MB (**fig. S6A**). Upon differentiation to MT, 822 genes exhibited a 2-fold (or more) increase in expression in WT cells as measured by RNA-seq while ~60% of these genes failed to properly induce in HDGF2 KO cells (**Figs. 4C & S6B-D**). Importantly, HDGF2 binding increased on these genes upon differentiation of WT cells, but not on genes that exhibited normal induction in the HDGF2 KO cells (**Fig. 4D**). In contrast, LEDGF exhibited a decrease in its global binding upon differentiation (**Fig. 4B & 4D**). An example of this phenomenon is observed at the myosin heavy chain (MHC) cluster that contains genes induced during differentiation to MT (**Fig. 4E**). Upon differentiation, HDGF2 levels increase on the induced genes within the cluster whereas LEDGF binding decreased uniformly across the entire cluster. SPT16 binding was undetectable on this cluster in both the MB and MT. The induced expression of these genes was dependent on HDGF2 as their induction was not detected by RNA-seq in HDGF2 KO MT as compared to WT MT (**Fig. 4E, bottom tracks**). These data strongly suggest that in myotubes, which contain undetectable levels of FACT and low levels of LEDGF, the binding of HDGF2 to chromatin is required for induction of its target genes.

We observed that most genes being expressed in both the myoblast and myotube were enriched in HDGF2 at both stages. An example of one such highly expressed gene (NES) that is dependent on HDGF2 is shown in Figure **4F**. However, RNA-seq analysis revealed many such genes that are not dependent on HDGF2 (**figs. S6B-C**). Therefore, the low levels of LEDGF in myotubes as well as additional factors that possess FACT-like activity likely exist for continued expression of these genes. One such additional factor may be NDF, which was recently identified in *Drosophila* embryo extract through a screen for factors that stimulate acetylation of nucleosomes *in vitro* by the acetyltransferase p300 (*34*). Notably, NDF (CG4747) and the fly homolog of LEDGF/HDGF2 (CG7946) were originally identified together by ChIP-MS as proteins that enriched with actively transcribed chromatin containing H3K36me3 (*29*).

Similar to 293T cells, we observed an opposition between H3K27me3 with H3K36me2, LEDGF and HDGF2 in MB (**Fig 2C-D & Figs. 4E-G**). Interestingly, upon differentiation to MT, a silent gene (C20orf166) in the MB located within an H3K27me3 domain, is induced and concurrently we observed a loss in H3K27me3 with an accumulation of HDGF2 (**Fig. 4G**). The expression of this gene was dependent on HDGF2, suggesting an interplay between HDGF2 and H3K27me3 domains during differentiation.

To better understand how HDGF2 affects transcription in cells, we employed PRO-Seq in the WT and HDGF2 KO myoblast cells. PRO-seq precisely maps the level of engaged RNAPII on genes as mature transcripts are washed away during the nuclear isolation process (*35*). Metagene profiles of PRO-seq data revealed an increased RNAPII occupancy within the promoter proximal region of many HDGF2 target genes in the HDGF2 KO as compared to WT cells (**Fig. 4H**). This data suggests that HDGF2 does not impact initiation, but rather the RNAPII transition to transcript elongation. Strikingly, the increased RNAPII peaks observed on genes in HDGF2 KO cells correlates with the first nucleosome position (**Fig. 4I**), indicating that HDGF2 helps relieve the nucleosome-induced block to transcription, consistent with our biochemical data (*36*). Notably this RNAPII pausing phenotype was observed in two independent HDGF2 KO myoblast cell lines, both of which could be rescued by stable lentiviral expression of HDGF2, demonstrating that the effect seen is due to HDGF2 elimination (**Fig. 4H-I’ right panels and S6E**).

In conclusion, we have identified two factors, LEDGF and HDGF2, as proteins which allow RNAPII to overcome the nucleosome-induced barrier to transcription elongation in differentiated cells that no longer express FACT (**Fig 4J**). During cellular differentiation histone modifications reorganize leading to distinct cells types with unique transcriptional profiles. In light of our findings, we propose that these proteins reorganize with histone modifications to maintain chromatin in a transcriptional competent state, hence overriding the requirement for FACT and helping to sustain the unique transcriptional profiles of particular cell types.

---

**Figure S1.**
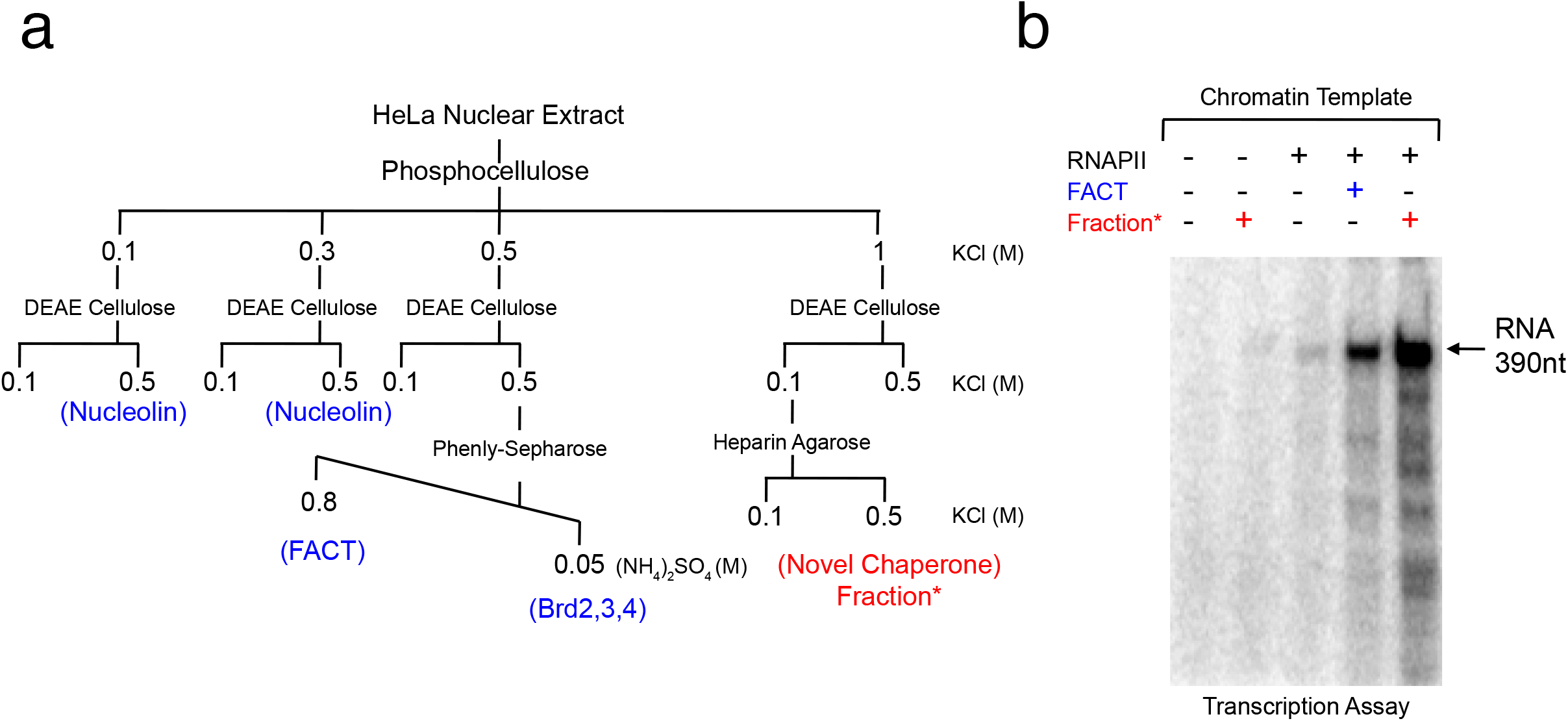
Identification of a Novel FACT-like chromatin transcription chaperone. **A.** Chromatographic fractionation scheme used to separate FACT-like activities present in Hela cell nuclear protein extracts and isolation of a novel FACT-like chaperone fraction. **B.** *In vitro* chromatin transcription assay used to screen for FACT-like activities. FACT or novel FACT-like chaperone fraction added as indicated.

**Figure S2.**
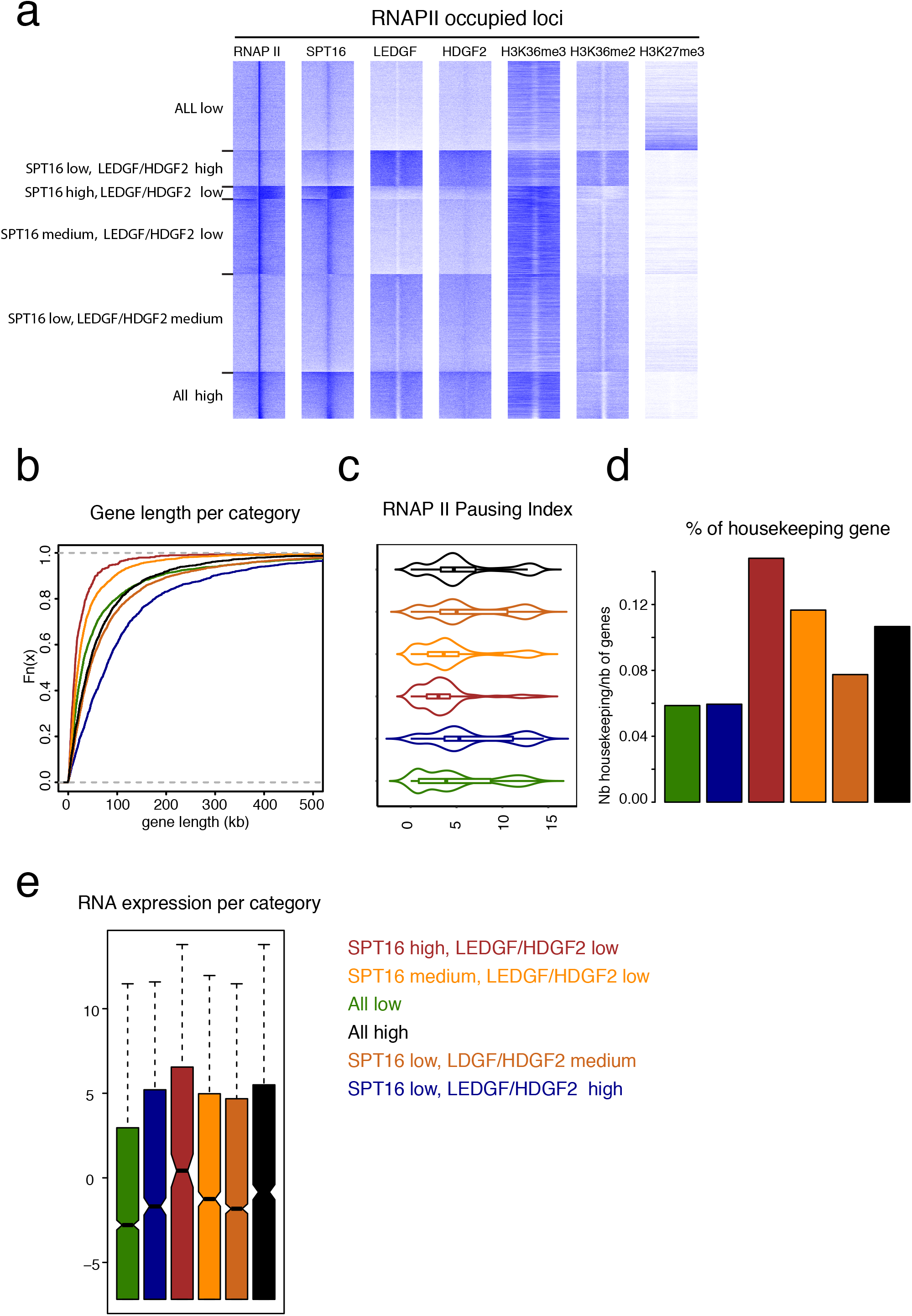
Bioinformatics analysis of genes within categories defined by levels of FACT, LEDGF and HDGF2 binding. **A.** Heatmaps showing six categories of genomic regions with RNAPII occupancy. Categories were defined by the levels of SPT16, LEDGF and HDGF2 (K-mean clustering) centered on RNAPII peaks and extending −10 Kb and +10 Kb. **B.** Cumulative plot of gene length per category. Y-axis indicates the cumulative sum of the gene length. X-axis indicates the gene length in kb. **C.** Violin plots showing the RNAPII pausing index per category. Pausing index in X-axis is defined by a ratio between RNAPII occupancy on promoters vs. gene bodies. **D.** Percentage of housekeeping genes in each category defined by TAU index (see methods). Y-axis indicates number (nb) of housekeeping genes normalized to the number of total genes. **E.** Boxplots of gene expression per categories indicated. Y-axis shows the log2 transcript per million (TPM) expression levels.

**Figure S3.**
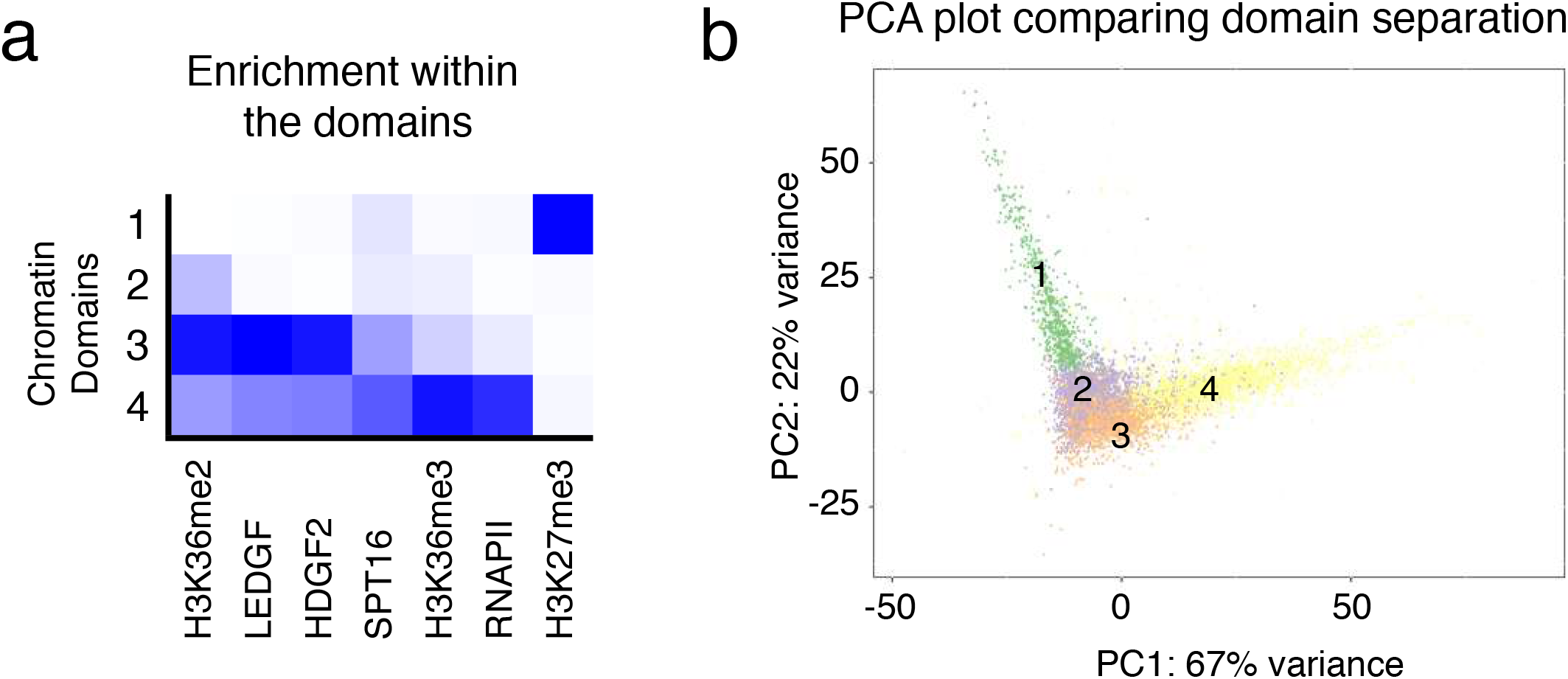
HDGF2/LEDGF co-localize with H3K36me2 and moderate levels of RNAPII excluding H3K27me3 domains. **A.** The 293T genome is partitioned into 4 chromatin domains based on the occupancy of the indicated factors. Markov modeling is used to determine the chromatin domains. The intensity of the blue color indicates the level of deposition/binding of each factor. **B.** Principal component analysis (PCA) plot representing partitioning of the loci defined by Markov modeling. Colors represent the 4 chromatin domains. PC1 represents 67% variance and PC2 represents 22% variance.

**Figure S4.**
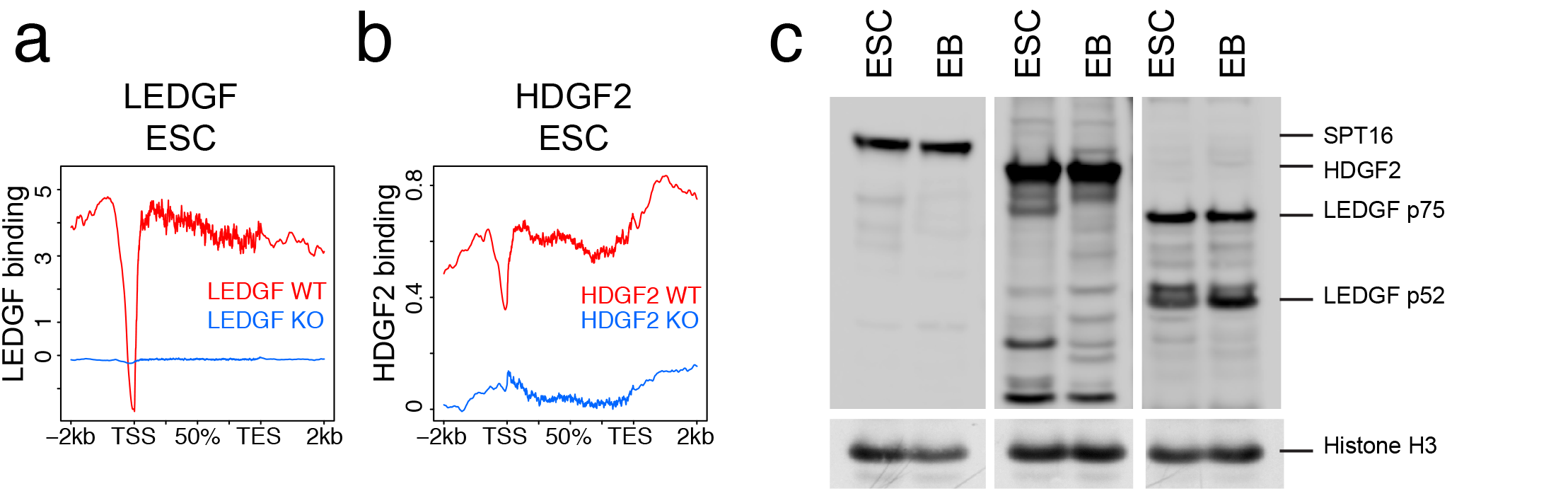
Validation of LEDGF and HDGF2 antibodies. **A., B.** Average density ChIP-seq profiles for LEDGF (**A**) and HDGF2 (**B**) with the indicated genotypes. Top 20% of the genes bound by the highest levels of either LEDGF or HDGF2 are used to generate the profiles along the TSS and TES extended by 2Kb from each site. **C.** Western blots of whole cell protein extracts from mESCs (ESC) differentiated to EBs (EB) using the antibodies as indicated.

**Figure S5.**
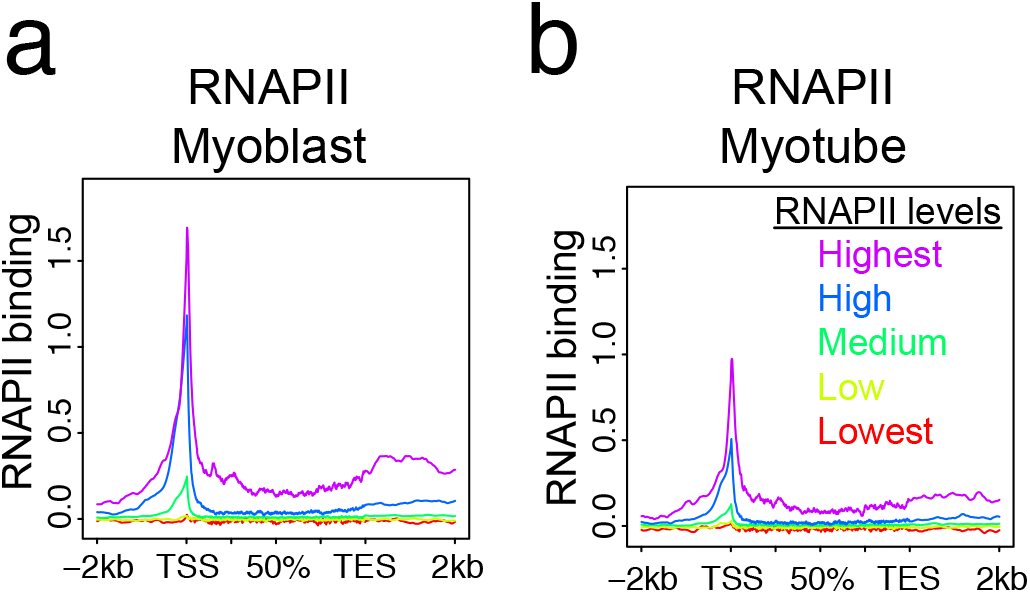
Genes in Myoblast and Myotube are categorized based on RNAPII occupancy. **A., B.** Average density ChIP-seq profiles for RNAPII in Myoblast (**A**) and Myotube (**B**) with the indicated gene categories determined by RNAPII levels: Highest (purple, top 5%), High (blue, top 5-20%), Medium (dark green, 20-80%), Low (bright green, bottom 5-20%) and Lowest (red, bottom 5%). Profiles are generated along the TSS and TES extended by 2kb from each site.

**Figure S6.**
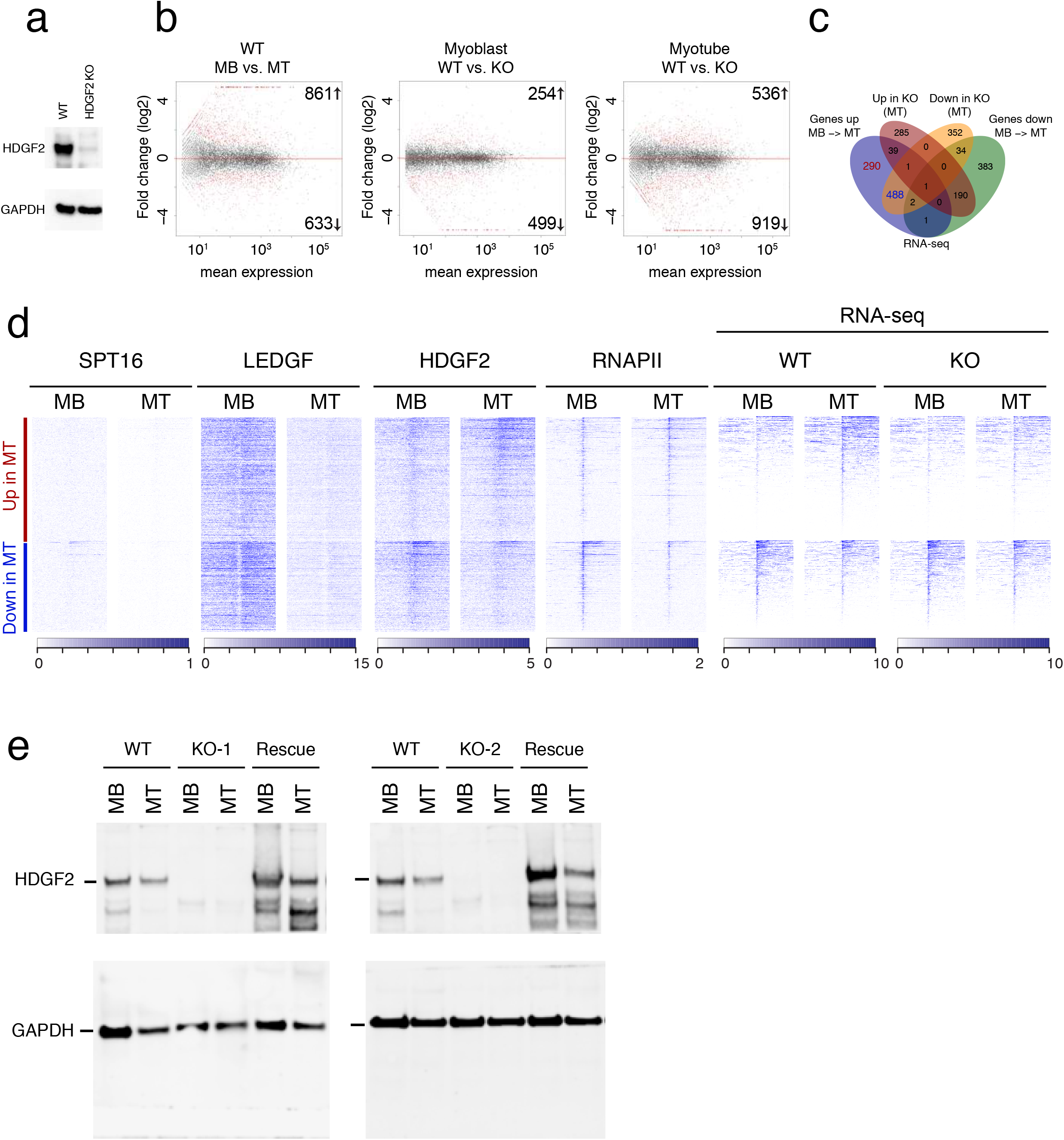
HDGF2 is required to activate Myotube specific genes. **A.** Western blots of whole cell protein extracts from WT or HDGF2 KO myoblasts using the antibodies as indicated. **B.** MA plots showing the number of differentially expressed genes (2-fold, BH-corrected p-value < 0.05) in WT cells (Myoblast vs. Myotube, left panel) and in WT vs. HDGF2 KO (Myoblast cells, middle panel) and (Myotube cells, right panel). **C.** Venn Diagram depicting the number of genes up-regulated 2-fold in MB to MT cells (blue) overlaid with genes down-regulated 2-fold in MB to MT cells (green), genes up-regulated 2-fold in HDGF2 KO MT cells (red) and genes down-regulated 2-fold in HDGF2 KO MT cells (yellow). **D.** Heatmaps of ChIP-seq read density of the indicated factors as well as RNA-seq read density on upregulated (red) and downregulated (blue) genes during Myoblast to Myotube differentiation. Heatmaps are centered on the TSS and extend −10 Kb and +10 Kb. Scales are indicated at the bottom of the Heatmaps. **E.** Western Blots of whole cell extracts from myoblasts and myotube cells. Derived cell lines indicated on top of the panels; WT cells, HDGF2 KO cells and HDGF2 KO cells rescued with a lentivirus that expresses HDGF2. Two independent HDGF2 KO cell lines loaded as indicated. Blots probed with antibodies as indicated.

## Acknowledgments

We thank Drs. L. Vales for her critical guidance and reading of the manuscript. We thank other members (past and present) of Reinberg lab for their discussion as the work was in progress. We are also grateful to D. Hernandez for technical assistance. Human myoblasts were obtained from as a generous gift from the Blau lab (Stanford University) and thanks to Stacy A. Marshall and Emily J. Rendleman for their help with PRO-Seq NGS (Northwestern University).

## Author contributions

G.L., O.O., N.D. and D.R. conceptualized and designed the study. G.L., O.O., R.G., H.O.K., C-H.L., J-R.Y., J.S. and Y.A. conducted the experiments. N.D. performed bioinformatics analyses. Y.A. and A.S. performed and analyzed the PRO-seq experiments. G. L., O.O., N.D. and D.R. wrote the manuscript.

## Data and materials availability

Genome sequencing results are available on GEO under accession number <u>GSE117155</u> and will be available publicly upon acceptance.

## Materials and Methods

### Purification and identification of LEDGF (PSIP1) from HeLa nuclear extract

HeLa nuclear protein extract (4g) was prepared as described in (*1*). Nuclear extract was dialyzed against BC100: BC buffer, pH 7.5 + 100mM KCL (20 mM Tris-HCl, 20 mM β-Mercaptoethanol, 0.2 mM PMSF, 0.2 mM EDTA, 10% glycerol (v/v) and 100 mM KCl). The number after BC denotes the salt concentration of the buffer. The extract was then loaded onto a phosphocellulose column and sequentially eluted with BC buffer containing 0.3, 0.5 and 1M KCl. The novel FACT-like activity eluted in the 1M KCL (BC1000) fraction. This fraction was dialyzed against BC100 and loaded on a DEAE Cellulose column and sequentially loaded with BC buffer containing 0.3, 0.5 and 1M KCl. The novel FACT-like activity did not bind the DEAE Cellulose and was collected in the flow through fraction (BC100). The flow through fraction was dialyzed against BA100: BA buffer, pH 7.5 + 100 mM NaCl (20 mM Hepes, 20 mM β-Mercaptoethanol, 0.2 mM PMSF, 0.2 mM EDTA, 10% glycerol (v/v) and 100 mM NaCl) and loaded onto a Heparin Agarose column. The column was washed with BA100 and eluted with BA600. The FACT-like activity eluted in BA600 fraction, which was then dialyzed against BC100 and loaded onto a Q-Sepharose column. The column was eluted sequentially with BC buffer containing 0.25 and 0.5 M KCl. The .25 M (BC250) fraction was dialyzed against BA100 and loaded onto a Heparin Agarose column. The Heparin Agarose column was washed with BA100 and eluted sequentially with 0.3, 0.5, 0.7 and 1 M NaCl in BA buffer. The novel activity eluted in the 0.7 (BC700) fraction. A portion of this fraction was then loaded on a Superdex-200 (gel filtration column) that was equilibrated and run in BC100. The activity eluted from the gel filtration column with a mass range between 150-75 KDa.

### Chromatin assembly and chromatin transcription assays

Chromatin assembly and chromatin transcription assays were performed as described in (*2, 3*).

### Recombinant LEDGF p75, LEDGF p55, HDGF2 and HDGF proteins

The cDNAs encoding the LEDGF p75, LEDGF p55, HDGF2 and HDGF proteins were cloned into the CBF vector (Addgene) at the SMA-1 site which when expressed makes a N-terminal FLAG Tagged fusion protein. The LEDGF p75, LEDGF p55, HDGF2 and HDGF proteins were produced and purified with the same protocol as used for Brd2, Brd3 and Brd4 described in (*2, 3*).

### Native FLAG-LEDGF and FLAG-HDGF2 ChIP-seq and ChIP-MS

Native FLAG-ChIPs for ChIP-seq and ChIP-MS were performed in 293T cells as described in (*4*). FLAG-LEDGF and FLAG-HDGF2 stable cell lines were produced using the pQCXIP retroviral vector (Clonetech).

### ChIP-seq

ChIP-seq experiments were performed as described before (*5*). In brief, nuclei was isolated from cells fixed with 1% Formaldehyde. Next, using a Diagenode Bioruptor, chromatin was fragmented into ~250bp. Chromatin immunoprecipitation was performed with the antibodies listed in below. Chromatin from *Drosophila* (1:100 ratio to the experimental chromatin) as well as *Drosophila* specific H2Av antibody was used as spike-in control in each sample. For ChIP-seq, libraries were prepared as described in (*6*) using 1-30 ng of immunoprecipitated DNA.

### RNA-seq

Total RNA from ESCs, EBs, 293Ts, MB and MTs was isolated with RNAeasy (Qiagen) and reverse transcribed using Superscript III and random hexamers (Life Technologies) to synthesize the 1st strand. Second strand was synthesized with dUTP to generate strand asymmetry using DNA Pol I (NEB, M0209L) and the E. coli ligase (Enzymatics, L6090L). RNA-seq libraries were constructed using the protocol described in (*6*).

### Antibodies

LEDGF (Proteintech) Rabbit Polyclonal, Cat # 25504-1-AP

HDGF2 (Proteintech) Rabbit Polyclonal, Cat # 15134-1-AP

SPT16 (Cell Signaling) Rabbit Monoclonal D712K, Cat # 12191

SSRP1 (Abcam) Mouse Monoclonal 10D7, Cat # ab26212

GAPDH (Cell Signaling) Rabbit Monoclonal 14C10, Cat # 2118

MHC/MYH1 (DSHB) Mouse Monoclonal MF 20

H3K27me3 (Cell Signaling) Rabbit Monoclonal C36B11, Cat # 9733

H3K36me2 (Cell Signaling) Rabbit Monoclonal C75H12, Cat # 2901

H3K36me3 (Abcam) Rabbit polyclonal, Cat # ab9050

RNAPII (Santa Cruz) Rabbit Polyclonal N-20, Cat # sc-899

Anti-FLAG (Sigma Aldrich) M2 Agarose gel, Mouse Monoclonal M2, Cat # A2220

Histone H3 (Abcam) Rabbit polyclonal, Cat # ab1791

H2Av (Active Motif) Rabbit Polyclonal, Cat # 39715

### Mouse ESC culture and differentiation

E14Tga2 (ATCC, CRL-1821) ESCs were grown in standard medium supplemented with LIF, 1 μM MEK1/2 inhibitor (PD0325901, Stemgent) and 3 μM GSK3 inhibitor (CHIR99021, Stemgent). For embryoid body differentiation, 600K mESCs were plated in suspension plates with medium containing DMEM (Life Technologies), 20% FBS, 1% NEAA, 1% Pen/Strep, 2 mM L-Glutamine, 100 mM Ascorbic acid (Sigma). After 5 days, EB colonies were collected for downstream applications.

### Myoblast cell culture and differentiation to Myotubes

Human myoblasts were obtained from as a generous gift from the Blau lab (Stanford University). Cells are seeded on gelatinized plates and grown in Ham’s F-10 media (Gibco/Thermo Fisher) supplemented with 15% fetal bovine serum (Atlanta Biologicals), 1mM sodium pyruvate (Sigma), 100ug/ml penicillin/streptomycin (Gibco/Thermo Fisher), 2.5ng/ml basic human fibroblast growth factor (Promega), 1X GlutaMax (Thermo Fisher) and 1uM Dexamethasone. Once 90% confluent the myoblasts are differentiated to myotubes by changing the media to DMEM (Gibco/Thermo Fisher) supplemented with 2% Horse Serum (Hyclone) and 100ug/ml penicillin/streptomycin (Gibco/Thermo Fisher). Differentiation proceeds for 6 days and the efficiency can be monitored with microscopy as myotubes are easily visually detected.

### 293T cell culture

293T cells were cultured as described in (*4*).

### CRISPR genome editing

gRNAs were designed using CRISPR design tool in https://benchling.com. All gRNAs below were cloned in pSpCas9(BB)-2A-GFP (Addgene plasmid #48138) or to a lentiviral vector pLKO.1-puro U6 sgRNA (Addgene plasmid #50920). The gRNAs were transfected into mESCs using Lipofectamine 2000 (Life Technologies) or infected into Myoblasts together with lentiCas9-eGFP (Addgene plasmid #63592). Single clones from GFP positive and/or puromycin resistant cells were genotyped and confirmed by sequencing.

### gRNAs

Following gRNAs are used to knockout human HDGF2:

hsHDGF2-gRNA-KO1: GCCACACGCCTTCAAGCCCG

hsHDGF2-gRNA-KO2: CACCGGCCCCATCTCCCGCG

Following gRNAs are used to knockout mouse HDGF2:

msHDGF2-gRNA-KO1: TACCATCCAGTTACTTAGGG

msHDGF2-gRNA-KO2: GCTCACCTGGAACTGGCCTA

Following gRNAs are used to knockout mouse LEDGF:

msLEDGF-gRNA-KO 1: TTTTCAATGGAGCTGATCTG

msLEDGF-gRNA-KO2: GGTCTCATTGGAACATCCTA

### HDGF2 rescue myoblast cell lines

Clonal HDGF2 KO myoblast cell lines were rescued with the lentivirus pLV-EF1a-IRES-Blast that had the HDGF2 cDNA inserted within the multiple cloning sites (BamH1 and EcoR1). Following infection, cells were selected for Blasticidin resistance. The HDGF2 cDNA was modified to make it resistant to the CRISPr gRNAs used to knockout HDGF2.

### Tissue expression

The levels of expression FACT subunits (SSRP1 and SPT16) shown in Figure 1B were obtained with the Expression Atlas (https://www.ebi.ac.uk/gxa/home) and ordered by hierarchical clustering.

### Data processing

All samples were sequenced with either an Illumina Hiseq or NextSeq. The adapters were removed by the sequencing facility and the quality of sequencing was assessed with FastQC: (http://www.bioinformatics.babraham.ac.uk/projects/fastqc/). Reads having less than 80% of quality scores above 25 were removed with NGSQCToolkit v2.3.3 (*7*) using the command IlluQC.pl -se $file_path N A -1 $percent -s $threshold -p $nb_cpu -o $out_path. Human hg19 and mouse mm10 from Illumina igenomes UCSC collection were used. ChIP-seq data were aligned with Bowtie (*8*) v1.0.0 allowing 3 mismatches and keeping uniquely aligned reads (bowtie -q -v 3 -p $nb_cpu -m 1 -k 1 --best --sam --seed 1 $bowtie_index_path “$file_path”). RNA-seq data were aligned with Tophat (*9*) v2.0.9 allowing 3 mismatches (tophat -N 3 -- bowtie1 -o $out_path -p $nb_cpu $bowtie_index_path $file_path). Sam ouputs were converted to Bam with Samtools (*10*) v1.0.6 (samtools view -S -b $file_path -o $out_path/$file_name.bam) and sorted with Picard tools v1.88 (http://broadinstitute.github.io/picard: java -jar SortSam.jar SO=coordinate I=$input_bam O=$output_bam). Data were further processed with Pasha (*11*) with the following parameters: WIGvs = TRUE, incrArtefactThrEvery = 7000000 for chip-seq and NA for rna-seq, elongationSize = NA for chip-seq and 0 for rna-seq. Input subtraction and scaling were performed with the function *normAndSubtractWIG* for 293TRex ChIP-seq. Data in mESC-embryoid bodies and myoblast/myotube differentiation system were spiked-in scaled with ChIPSeqSpike (*12*). Fixed steps wiggle files were converted to bigwigs with the script *wigToBigWig* available on the UCSC Genome Browser website (http://hgdownload.soe.ucsc.edu/admin/exe/).

### Venn diagrams

Signal was first detected with macs2 (*13*) (macs2 callpeak -t $bam_file_vector -c $input_file_vector -n $experiment_name --outdir $output_folder_nomodel_broad -f $format -g $genome_size -s $tag_size --nomodel --extsize $elongation_size --keep-dup $artefact_threshold --broad --broad-cutoff $qvalue). The parameters --extsize and --keep-dup were determined from the Pasha output log. Overlap was determined with the *findOverlapsOfPeaks* function of the ChIPpeakAnno (*14*) package (Figures 1A, 2B, 3E and 4C).

### Heatmaps

The heatmap of Figure S2A was obtained using a K-mean clustering on SPT16 and LEDGF with 6 groups using the Bioconductor package *seqplots* (*15*). Intervals are centered on the maximum Pol II signal -/+ 10 kb and correspond to a macs2 peak detection (31,408 peaks) with broad mode and q-value < 0.03 (see venn diagrams section for details). Other marks are plotted on the above defined loci.

### Correlation plot

Figure 2D is a spearman correlation on the top 20% bound loci composed of the union of H3K27me3 and H3K36me2 (24,289) peaks. The peaks were obtained with macs2 in broad mode with q-value < 0.04.

### Differential binding analysis

In mouse ESC to embryoid bodies differentiation system, the differential binding analysis (Figure 3C) was performed with DiffBind (*16*) on Refseq gene annotations. The percentages were calculated by computing the overlap of the different up-bound genes for each mark.

### Differential expression

Differentially expressed genes were obtained with DESeq2 (*17*) (Figure 3D-E, 4C & S6B).

### Metagene profiles

Metagene profiles (Figure 3G, Figure 4D) were performed with the bioconductor package seqplots. Mean, 95% confidence intervals and standard error are indicated on the profiles. The meta-profiles of Figures 4B and S5 were obtained with in-house scripts that divide gene mean values into Highest (purple, top 5%), High (blue, top 5-20%), Medium (dark green, 20-80%), Low (bright green, bottom 5-20%) and Lowest (red, bottom 5%) bound genes.

### RNA Pol II pausing

The violin plot of Figure S2C shows the pausing of Pol II for the genes present in the different categories defined on the heatmap of Figure S2A. The pausing index was calculated as being the ratio of the mean values at TSS-300bp+100bp and 50% of gene body to TES. The number of genes for the different categories in Figure S2A (top to bottom) are as follows; 3159, 1492, 487, 2002, 4288 and 2013 respectively.

### Housekeeping vs tissue specificity expression

In Figure S2D, the house keeping feature of each gene of the different categories defined in Figure S2A was calculated using the *tau* metrics (*18*) using a threshold of 0.15 on the tissues as described in (*19*). Similar results were found using (*20*) and (*21*) (data not shown).

### Hidden markov model

The markov model of Figure S3A was calculated using ChromHMM v1.12 (*22*) with 4 states and a window size of 80 kb.

### PRO-seq library preparation and sequence alignment

PRO-seq was performed according to the protocol (*23*) with minor modifications. In the nuclear run-on reaction using myoblast with spike-in *drosophila* S2, all 4 biotinylated nucleotides were used at 25 μM each final concentration. RppH (NEB) was used to remove the 5’ RNA cap. DNA Libraries were size selected by AMPure XP beads (Beckman Coulter) and sequenced on a NextSeq 500. Adaptors were removed from raw reads by cutadapt 1.14 (*24*). Reads were trimmed from the 3’ end with removing low quality bases using Trimmomatic 0.33 (*25*) requiring a read length of 16–36 bp. Reads derived from ribosomal RNA were filtered out by mapping reads on human and fly ribosomal DNA. The remaining reads were mapped on human hg19 or fly dm3 genome using Bowtie 1.1.2 (*8*) with options -m 1 -v 2. The 5’ ends of the reads were taken using bedtools genomecov 2.25 (*26*) with options -strand -bg -5 and strands of reads were then swapped. Read counts were normalized by spike-in aligned reads. Normalized read counts were divided by total aligned million reads of wild-type sample. Metagene profiles of PRO-seq were generated by deepTools 3.0.0. computeMatrix and plotProfile (*27*).

